# Comparative Analysis of In Vitro Responses and Regeneration Between Diverse Bioenergy Sorghum Genotypes

**DOI:** 10.1101/861328

**Authors:** Barry S. Flinn, Savanah Dale, Andrew Disharoon, Stephen Kresovich

## Abstract

Sorghum has been considered a recalcitrant plant in vitro, and suffers from a lack of regeneration protocols that function broadly and efficiently across a range of genotypes. This study was initiated to identify differential genotype-in vitro protocol responses across a range of bioenergy sorghum bioenergy parental lines, in order to characterize response profiles for use in future genetic studies. Seven bioenergy sorghum genotypes were compared, along with the common grain sorghum genotype Tx430, for their in vitro regeneration responses using two different in vitro protocols, LG and WU. All genotypes displayed some level of response during in vitro culture with both protocols. Distinct genotype-protocol responses were observed, with the WU protocol significantly better for plantlet regeneration. All bioenergy genotypes, with the exception of Chinese Amber, performed as well, if not better than Tx430, with Rio and PI329311 the top regenerating lines. Genotypes displayed protocol-dependent, differential phenolic exudation responses, as indicated by medium browning. During the callus induction phase, genotypes prone to medium browning exhibited a response on WU medium which was either equal or greater than on LG medium, with Pink Kafir and PI329311 the most prone to medium browning. Genotype- and protocol-dependent albino plantlet regeneration was also noted, with three of the bioenergy genotypes showing albino plantlet regeneration. Grassl, Rio and Pink Kafir were susceptible to albino plantlet regeneration, with the response strongly associated with the WU protocol. Pink Kafir displayed the highest albino formation, with close to 25% of regenerating explants forming albino plantlets.

## Introduction

Sorghum [*Sorghum bicolor* (L.) Moench] ranks fifth of the major grain crops in production, area harvested, and yield worldwide (FAOSTAT Database 2017), and more than 300 million people use it as a staple food, particularly in developing semiarid tropical regions (Kebede et al. 2001). In addition to its use as a human food source, sorghum is also used for animal feed (Mabelebele et al. 2018), brewery or bio-functional malted beverages (Garzón and Drago 2018), building materials (Khazaeian et al. 2015), as a source of sweet syrup/juice (Asikin et al. 2018), a source of bioactive metabolites (Vanamala et al. 2018), and for bioenergy (Rooney et al. 2007). It is a hardy crop, able to withstand both drought and flooding conditions, as well as produce high yields, making it a model crop for agricultural adaptation to climate change and human population growth (Paterson et al. 2009).

As with most crops of value, there are concerns associated with sorghum production, including losses due to abiotic and biotic stress pressure, as well as the desire to enhance composition and increase yield, indicating that a variety of targets exist for trait improvement. Improvement efforts have utilized breeding, coupled with approaches such as QTL identification and mapping, molecular marker identification and genome wide association studies, with the goal of improving germplasm while also gaining an understanding of the genetic loci/genes/allelic variation contributing to the various traits (Chopra et al. 2015, Ali et al. 2016, Boyles et al. 2016, Brenton et al. 2016, Boyles et al. 2017, Mofokeng et al. 2017, Disasa et al. 2018, Boyles et al. 2019, Mace et al. 2019).

Plant tissue culture, coupled with in vitro regeneration, can be integrated into sorghum trait improvement programs, providing opportunities for trait modification without the temporal restrictions of traditional breeding, through somaclonal mutagenesis (Bhaskaran et al. 1987), the rapid development of homozygous lines by anther culture to create haploid plants, followed by chromosome doubling (Kumaravadivel and Rangasamy 1994), and the development of improved germplasm through the creation of transgenic (Reddy et al. 2015) or genome-edited (Char et al. 2019) plants. Sorghum has been the subject of several tissue culture studies, in which a variety of protocols have been used with a relatively small number of genotypes. These studies have shown that sorghum in vitro responsiveness displays genotype-dependency (Kaeppler and Pedersen 1997, Liu et al. 2015, Omer et al. 2018), with sorghum considered to be relatively recalcitrant to in vitro manipulation, due to factors like genotype-dependent phenolic production and tissue browning. Efforts to improve this recalcitrance have used different tissue culture media modifications to improve cell survival (Elkonin et al. 1995, Liu et al. 2015, Dreger et al. 2019) and subsequent regeneration, as well as the overexpression of *Baby Boom* and *Wuschel* genes to stimulate the embryogenic potential of explant cells, and improve regeneration (Lowe et al. 2016). An understanding of the genetics associated with tissue culture/in vitro responsiveness would facilitate breeding for improved regenerability/transformability, but also provide for a greater understanding of sorghum developmental processes. Furthermore, as regenerability is associated with organ formation, a better understanding of the genetics of in vitro responsiveness and regeneration could improve our understanding of the genetic loci/genes associated with the improved resource allocation required for new or replacement organ development, which would be of relevance to yield and performance enhancement (Arikita et al. 2013). Several studies have used QTL analysis and GWAS to identify genetic loci and candidate genes involved in tissue culture responsiveness and regeneration in a limited number of plant types (Tyagi et al. 2010, Begheyn et al. 2018, Ma et al. 2018), but an understanding of the overall genes and pathways associated with this process remains to be resolved, especially for recalcitrant sorghum.

Our lab has been focused on the use of genetic diversity to improve a wide variety of sorghum traits, with major interests in bioenergy, sweet, grain and forage types. A key resource of interest to the sorghum and overall plant research community was developed, the Bioenergy Association Panel (BAP), comprising genetically diverse cellulosic and sweet sorghum germplasm (Brenton et al. 2016). A subset of the BAP was used as parents to create Nested Association Mapping (NAM) populations, where nine NAM populations were developed. All NAM parental lines and the population individuals have been characterized through genotyping-by-sequencing (GBS), with whole genome sequencing of the NAM parental lines planned for the future. These NAM parents and the resultant populations provide a new resource to explore the genetics associated with a variety of sorghum traits. To initiate a characterization of the NAM parental lines for in vitro responsiveness and regeneration, their performance was assessed during culture under two different in vitro protocols, and compared against genotype Tx430, a grain type sorghum commonly used for in vitro studies. The goal was to identify differential genotype-protocol responses that could be used for future genetic studies.

## Materials and methods

### Plant materials

A variety of sorghum genotypes, representing various types, races and location of origin were used in this study (Table 1). Mature seeds were planted in 3 gallon pots containing Sungro Fafard® germination mix (Agawam, MA, USA) fertilized with the appropriate dose of Scotts Osmocote Classic 14-14-14 pellets (Marysville, OH, USA), and placed in the Clemson University Biosystems Research Complex greenhouse for germination and growth. Plants were checked daily and watered as necessary. At the onset of pollen production, each floral spike was bagged, and allowed to develop for 14 days. After 14 days, developing panicles were excised and removed to the laboratory. Immature seeds were removed from the middle third of each panicle, placed into sterile 50 ml conical tubes and sterilized with 45 ml of 20% concentrated commercial bleach (8.25% sodium hypochlorite active ingredient) containing two drops of Tween 20, with sterilization for 25 min at room temperature on an orbital shaker (220 rpm). In a sterile tissue culture hood, the bleach solution was removed, seeds were rinsed five times each with 45 ml of sterile RO water, and then transferred to a sterile petri dish for storage while embryo excisions took place.

**Table 1.**
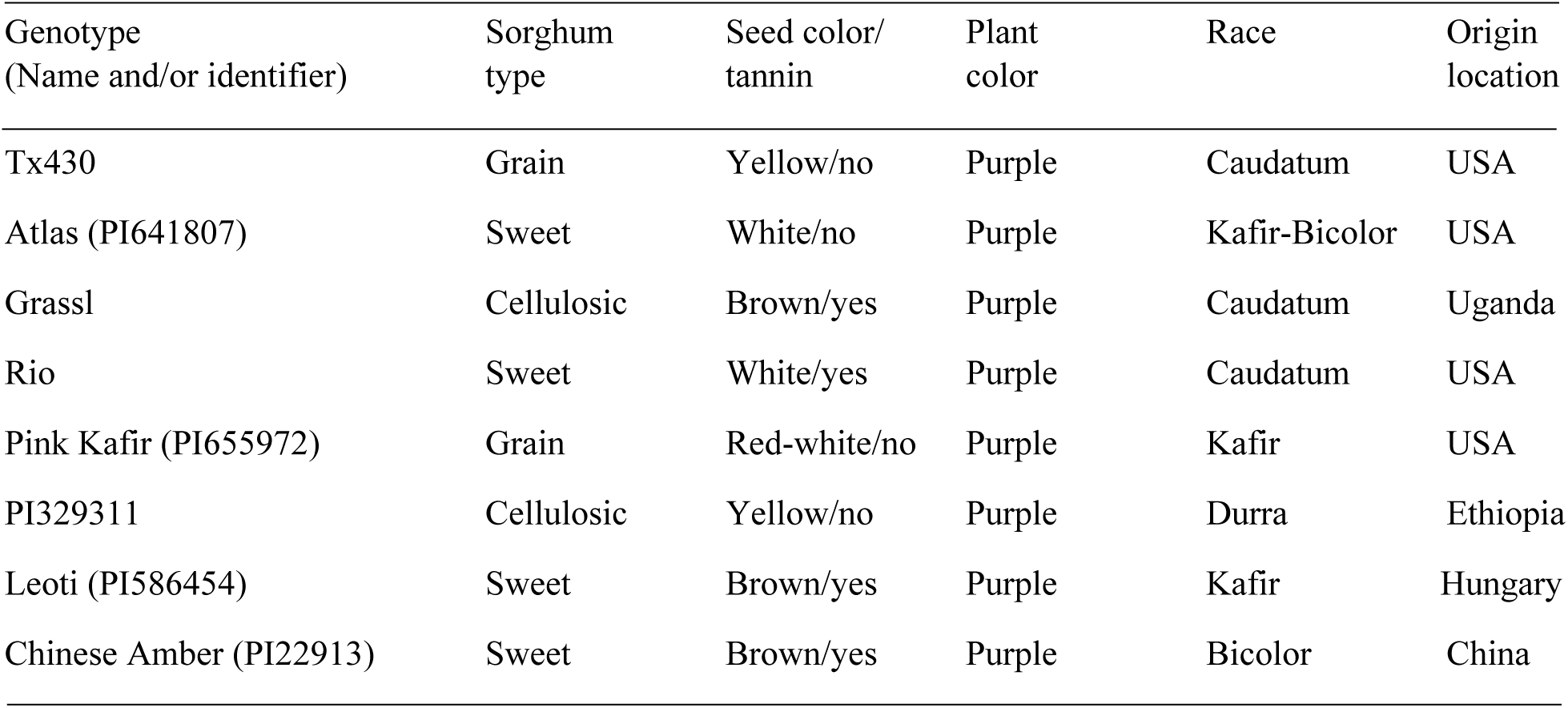
Sorghum genotypes used in this study.

### Tissue culture media

Two different tissue culture protocols, using Murashige and Skoog basal salts (Murashige and Skoog 1962), were tested with all genotypes and were adapted from Liu and Godwin (2012) and Wu et al. (2014). All media components were obtained from Phytotechnology Laboratories (Lenexa KS, USA), except for the gelling agents (agar and Phytagel^TM^), which were obtained from Sigma-Aldrich (St. Louis, MO, USA). As both original published papers were focused on transformation, in this current work, no selection agents were included in the media preparations. The LG protocol used Callus Induction Medium (CIM) and Regeneration Medium (RM) as described (Liu and Godwin 2012). The WU protocol used DBC3 and PHI-XM media (Wu et al. 2014) as CIM and RM, respectively. All compounds for both CIM and WU RM were added prior to pH adjustment and autoclaving for 15 min at 121°C, with the exception of copper sulfate, which was added as a sterile solution to pre-cooled (55°C) media. During LG RM preparation, growth regulators were added post-autoclaving to pre-cooled (55°C) medium. Briefly, to prepare a final 1 L of medium, 960 ml was prepared, and prior to the addition of agar, growth regulators and copper sulfate, a 160 ml aliquot was removed. The remaining 800 ml aliquot was pH adjusted, the agar added, the solution autoclaved as above, and then cooled to 55°C in a water bath. To the 160 ml aliquot, all growth regulators were added, the solution brought to 200 ml with RO water, pH adjusted to 5.7, and then filter sterilized using a 250 ml 0.2 μm filter unit. This 200 ml was added to the 800 ml pre-cooled autoclaved fraction, followed by the addition of the sterile copper sulfate solution. CIM and RM were dispensed as 20 ml aliquots into sterile 100 x 15 mm petri plates, air dried for 25 min in the hood, and then stored in the dark at 6°C until use. Prior to use, plates were allowed to equilibrate to room temperature in the dark for several hours.

Following culture on each RM tissue culture regime, explants were transferred to Elongation and Rooting Medium (ERM) - ½ MS salts, ½ MS vitamins, sucrose (30 g/L), Phytagel^TM^ (3 g/L), pH 5.8. In this case, the autoclaved medium was dispensed as 50 ml aliquots into sterile, Magenta GA-7 vessels (Sigma-Aldrich, St. Louis, MO, USA), allowed to air dry for 25 min in the hood prior to vessel closure, and then stored at 6°C until use. Prior to use, vessels were allowed to equilibrate to room temperature in the dark for several hours.

### Explant culture

Immature embryos were excised aseptically under a dissecting microscope, and placed scutellum side up on the appropriate CIM, with 12 embryos per plate and approximately 60 embryos per medium per genotype per replicate, with 3 replicates per medium per genotype. Plates were wrapped in foil and incubated in the dark for two weeks for callus induction. Following the two weeks, all explants were transferred to their respective regeneration medium (RM), retaining the 12 explants per plate arrangement. All explants were cultured for a total of four weeks on RM, with subculture to fresh RM after the initial two weeks. The LG protocol explants were exposed to a 16 h light/8 h dark regime for the entire four-week period. The WU protocol explants were wrapped in foil and cultured in the dark for the first two weeks, and then cultured in the light/dark regime described above for the remaining two weeks. Explants were cultured in a VWR incubator with a 27°C light (16 h) /20°C dark (8 h) cycle, and a light intensity of 70 µmol m^-2^ s^-1^.

### Tissue culture data collection

All explants, unless they exhibited fungal/bacterial contamination, were transferred to the appropriate media for the complete eight-week in vitro culture scheme. Digital photos were taken of all explants for each genotype/media replication at the end of culture on CIM, RM and ERM. The photos taken after culture on CIM were used for quantification of medium browning, as described below. Following culture on RM, explants were qualitatively assessed overall for their degree of callus proliferation, as well as for medium browning. A callus proliferation index (+, ++, +++, ++++) based on qualitative, visual assessment of callus for all explants relative to initial embryo explants per treatment was made for genotype/media combinations. Similarly, a medium browning index (+, ++, +++, ++++) based on qualitative, visual assessment of medium browning for all explants relative to initial embryo explants per treatment was made for genotype/media combinations. Upon completion of the eight-week culture scheme, explants were assessed for the number of surviving, non-necrotic explants, and the number of surviving explants that exhibited plantlet regeneration, with the information used to generate percentage response. For each explant, independent and rooted plantlets could easily be removed from explant callus and were counted, to provide a determination of the number of regenerants per responding explant. Mean values, standard deviations and standard errors were calculated.

### Image analysis

Digital color images of petri plates taken for all genotypes following two weeks of culture on CIM were quantified using the Fiji image-analysis software platform (Schindelin et al. 2012). Color images were converted to 8-bit black and white images, and the oval/elliptical tool used to outline each plate, followed by a determination of the total plate area using the area measurement function. The free hand line tool was used to outline any visibly darkened region of media around each individual explant per plate, and the total area of browning determined using the area measurement function. The total media browning area per plate was determined by adding the browning area measurements, and then expressing this combined area as a percentage of the total petri plate area.

### Statistical analysis

For experiments, data comparisons were made between all genotypes within a tissue culture protocol (all genotypes on LG, all genotypes on WU) and within each individual genotype on LG and WU. Data was subjected to one-way ANOVA, followed by Tukey’s HSD Post-hoc Test. Data which showed a probability of P=0.05 or less were considered significantly different.

## Results

Bioenergy sorghum trait improvement is a current focus of several labs, with tissue culture representing one tool for use in this effort. The grain sorghum genotype Tx430 has been commonly and successfully used in transformation and regeneration studies. Therefore, this genotype was used as a baseline for comparative purposes to characterize the in vitro responses of several diverse bioenergy sorghum genotypes used as parents to create NAM populations.

The general tissue culture steps for both in vitro protocols (LG and WU) are shown using Tx430 as the representative genotype (Fig. 1). Freshly excised 14 DAP embryos were placed on LG or WU Callus Induction Medium (CIM) and cultured in the dark for two weeks. During this period, callus induction and growth took place from the explant scutellum, with the formation of white and cream-colored calli, or in some cases, browning of the explant and/or medium. After the CIM phase, all explants were transferred to the respective LG or WU Regeneration Medium (RM), to allow additional callus proliferation and embryo/plantlet formation. Explants were cultured on RM for a total of four weeks, with subculture to fresh medium after two weeks. Following culture on RM, explants on both media exhibited distinct yellow, green, white and/or browning regions. Distinct morphological structures were observed arising from proliferating calli, with explant calli on both LG and WU RM forming shoots. The small shoots were more elongated on LG medium compared to those on WU medium, and generally more adventitious root growth from the calli cultured on LG RM. All explants were then transferred to Elongation and Rooting Medium (ERM) for a further two weeks under a 16 h photoperiod, which facilitated substantial shoot elongation and rooting, producing well-defined rooted plantlets that could easily be plucked from the associated explant callus. Regenerated plantlets were viable and could be transferred ex vitro into soil, acclimatized and grown in the greenhouse (data not shown). As our goal was to assess the overall in vitro responses of explants representing the various bioenergy genotypes for the two (LG, WU) protocols, rather than optimize the regeneration responses/yield of regenerants, we did not separate out regenerating and non-regenerating sectors, but kept each explant as intact as possible throughout all subcultures.

**Figure 1.**
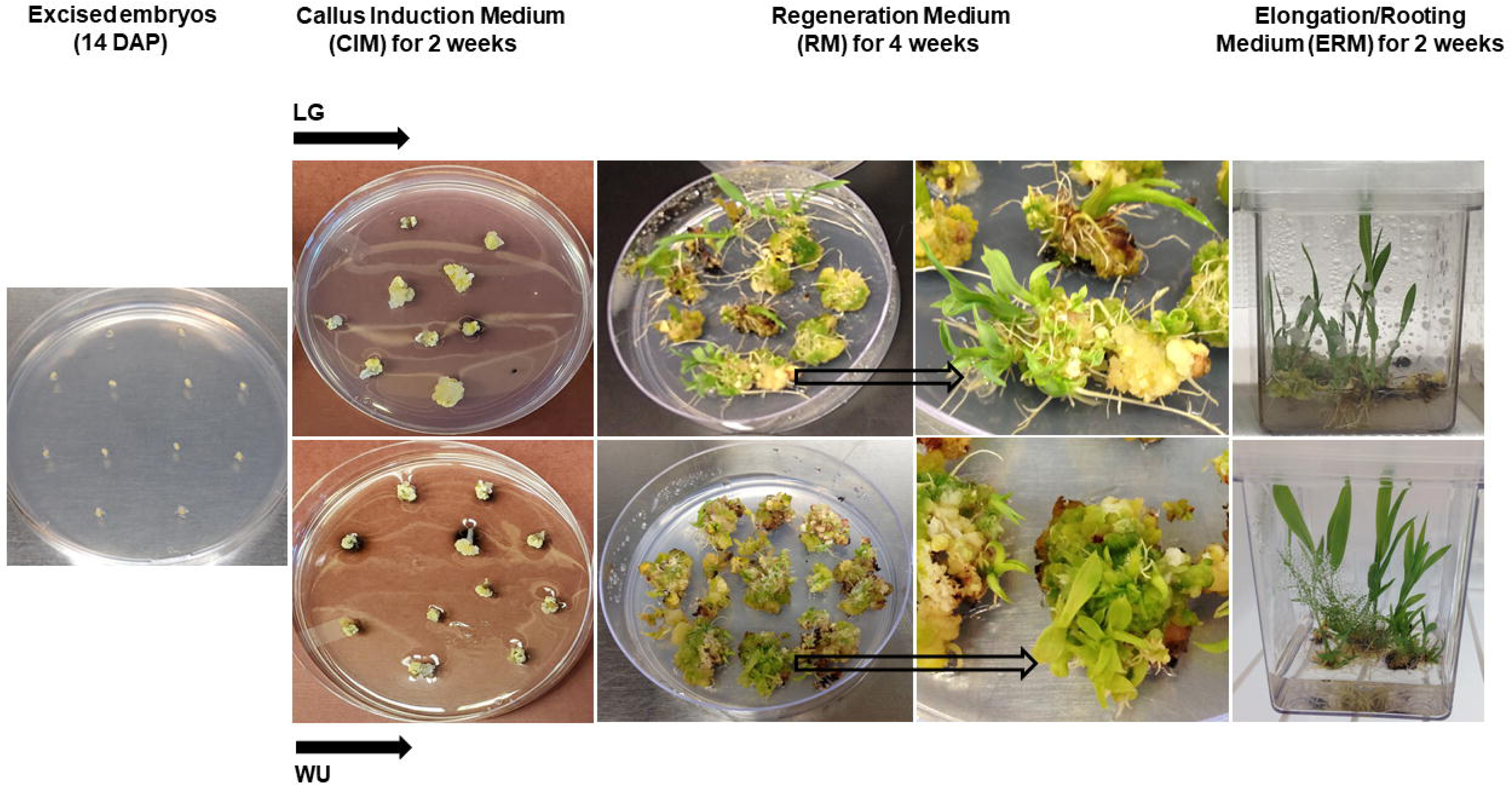
The in vitro tissue culture steps used for LG and WU protocols, leading to plantlet regeneration, with Tx430 shown as the representative genotype. Arrows denote individual explants enlarged in the right RM panel photo.

### CIM Browning Responses

As we followed the in vitro pathway outlined above for Tx430 and the seven genetically diverse NAM parental genotypes, distinct differences were noted for genotype and media combinations during the two weeks of culture on CIM. A common response phenotype was phenolic secretion by explants into the medium, with subsequent medium browning (Fig. 2). A comparison of explants cultured on LG CIM, exhibited very little medium browning with Tx430 and Leoti. In contrast, Pink Kafir exhibited maximal medium browning, while the other genotypes were intermediate in medium browning. On WU CIM, Tx430 and Leoti also showed minimal browning response, while Pink Kafir and PI329311 exhibited the highest browning levels. Comparisons within each genotype across LG and WU CIM revealed a significant increase in browning on WU CIM for Tx430 and PI329311. We never observed a higher level of browning for any genotype on LG CIM when compared against WU CIM, indicating that the WU protocol was more supportive of browning in genotypes prone to phenolic exudation.

**Figure 2.**
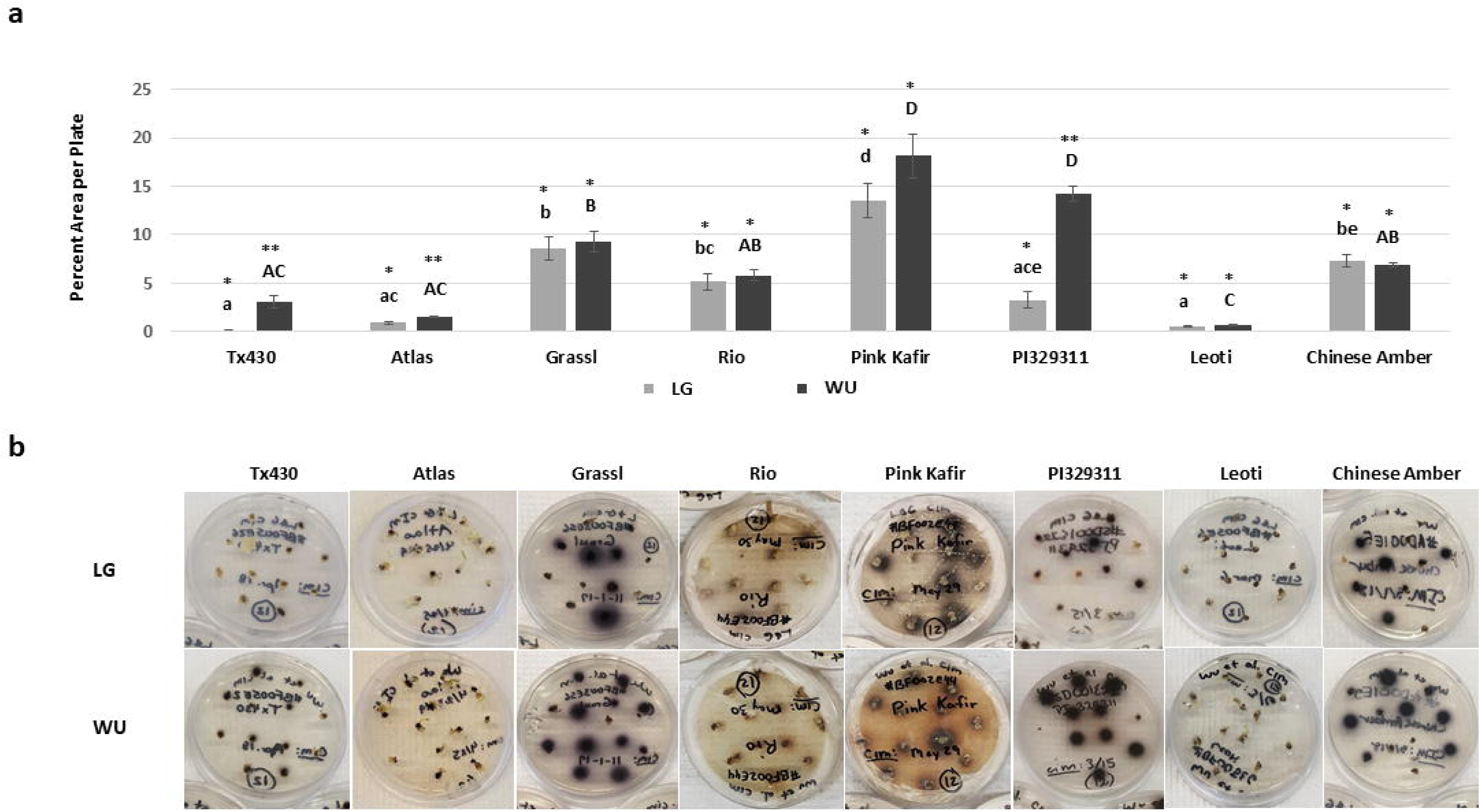
Impact of LG and WU CIM on medium browning from immature embryo explants of diverse sorghum genotypes, following two weeks of culture on CIM. a) Digital images were analyzed as described in the Materials and Methods, and plotted as the mean area per petri plate (± SE) exhibiting medium browning. b) Representative photos of petri plates for each genotype and protocol combination illustrating the levels of medium browning. Data shown in (a) were analyzed by one-way ANOVA, and tested for significance using Tukey’s HSD Post-hoc Test. For comparisons across genotypes for LG CIM, those with the same lowercase letter are not significantly different at P≤0.05. For comparisons across genotypes for WU CIM, those with the same uppercase letter are not significantly different at P≤0.05. For comparisons across the two media within a genotype, those with the same asterisk pattern are not significantly different at P≤0.05.

### RM Responses

Explants were transferred from CIM to RM and cultured for four weeks. During culture on RM, explants often exhibited continued callus proliferation and an overall increase in size and mass on both media, although there was an impact of genotype and media combination on observed responses (Fig. 3). A visual qualitative proliferation index (Table 2) was used to provide a measure of callus proliferation relative to the initial excised embryo placed into culture. On LG RM, Tx430 and PI329311 displayed the greatest degree of callus proliferation, while Rio, Leoti and Chinese Amber were the poorest, with Chinese Amber the worst. On WU RM, Tx430, Pink Kafir and PI329311 were the most proliferative, while Leoti and Chinese Amber were again the poorest in callus proliferation. Comparisons within each genotype across LG and WU RM revealed that callus proliferation for Tx430, PI329311, Leoti and Chinese Amber were similar regardless of culture of LG or WU RM, while Grassl, Rio and Pink Kafir proliferated better on WU RM than on LG RM. Again, we never observed a better response within each genotype on LG RM when compared against WU RM. The poor performance of Chinese Amber explants is most likely a reflection of the extreme browning of the explants themselves, as this genotype exhibited a very pronounced response in which the explants became very soft and mushy.

**Figure 3.**
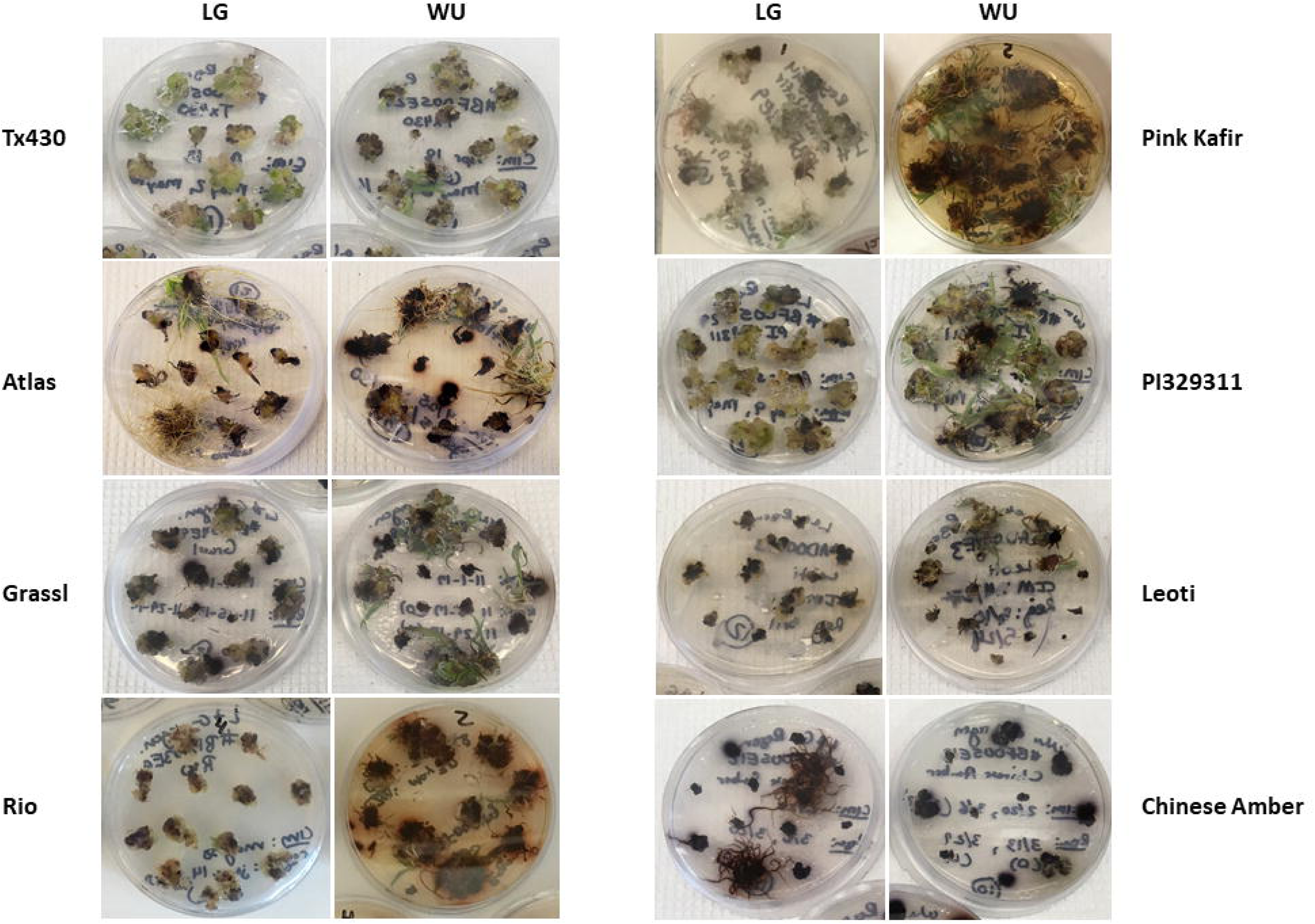
Impact of LG and WU RM on explant phenotype for the various genotypes, following four weeks of culture on RM. Representative photos of petri plates for each genotype and protocol combination illustrating the levels of callus proliferation and medium browning.

**Table 2.** Impact of genotype and protocol on callus proliferation and medium browning on RM.

| Genotype | Protocol | Proliferation index | Medium browning |
| --- | --- | --- | --- |
| Tx430 | LG | ++++ | - |
|  | WU | ++++ | + |
| Atlas | LG | ++ | + |
|  | WU | +++ | ++ |
| Grassl | LG | ++ | +++ |
|  | WU | +++ | +++ |
| Rio | LG | ++ | + |
|  | WU | +++ | ++++ |
| Pink Kafir | LG | +++ | + |
|  | WU | ++++ | ++++ |
| PI329311 | LG | ++++ | + |
|  | WU | ++++ | ++ |
| Leoti | LG | + | ++ |
|  | WU | + | ++ |
| Chinese Amber | LG | + | +++ |
|  | WU | + | +++ |

In addition to callus proliferation, we also observed some medium browning responses during culture on RM (Fig. 3). While most genotypes exhibited minimal browning across LG RM and WU RM, the most intense medium browning was observed with Rio and Pink Kafir, with the response being minimal on LG RM, but extremely intense on WU RM. We noted that Grassl and PI329311, which displayed significant browning on WU CIM, exhibited less browning on WU RM. As noted above for CIM culture, within each genotype comparison, we never observed a more intense browning response on LG when compared against WU; browning responses were either similar for a genotype on both RM media, or more intense on WU RM.

### Performance following ERM culture

Following four weeks on RM, all explants were transferred to a growth regulator-free elongation and rooting medium (ERM) to allow plantlet elongation and rooting. After two weeks on ERM, explants were assessed for final performance metrics. Analysis of the overall survival rate (Fig. 4a) revealed no significant difference between any genotype cultured using the LG protocol. While Tx430 and PI329311 displayed the highest survival rates (100%) and Rio the lowest (50%), there was sufficient variability in survival across experiments to confer no significant differences. Using the WU protocol, we again noted that the highest survival rates were with PI329311 (99%) and Tx430 (87%), and the lowest with Chinese Amber (37%). Most of the genotypes were not significantly different from each other with respect to survival, except for PI329311, which was significantly greater than Chinese Amber. No significant difference in explant survival was observed within each genotype between LG and WU protocols.

**Figure 4.**
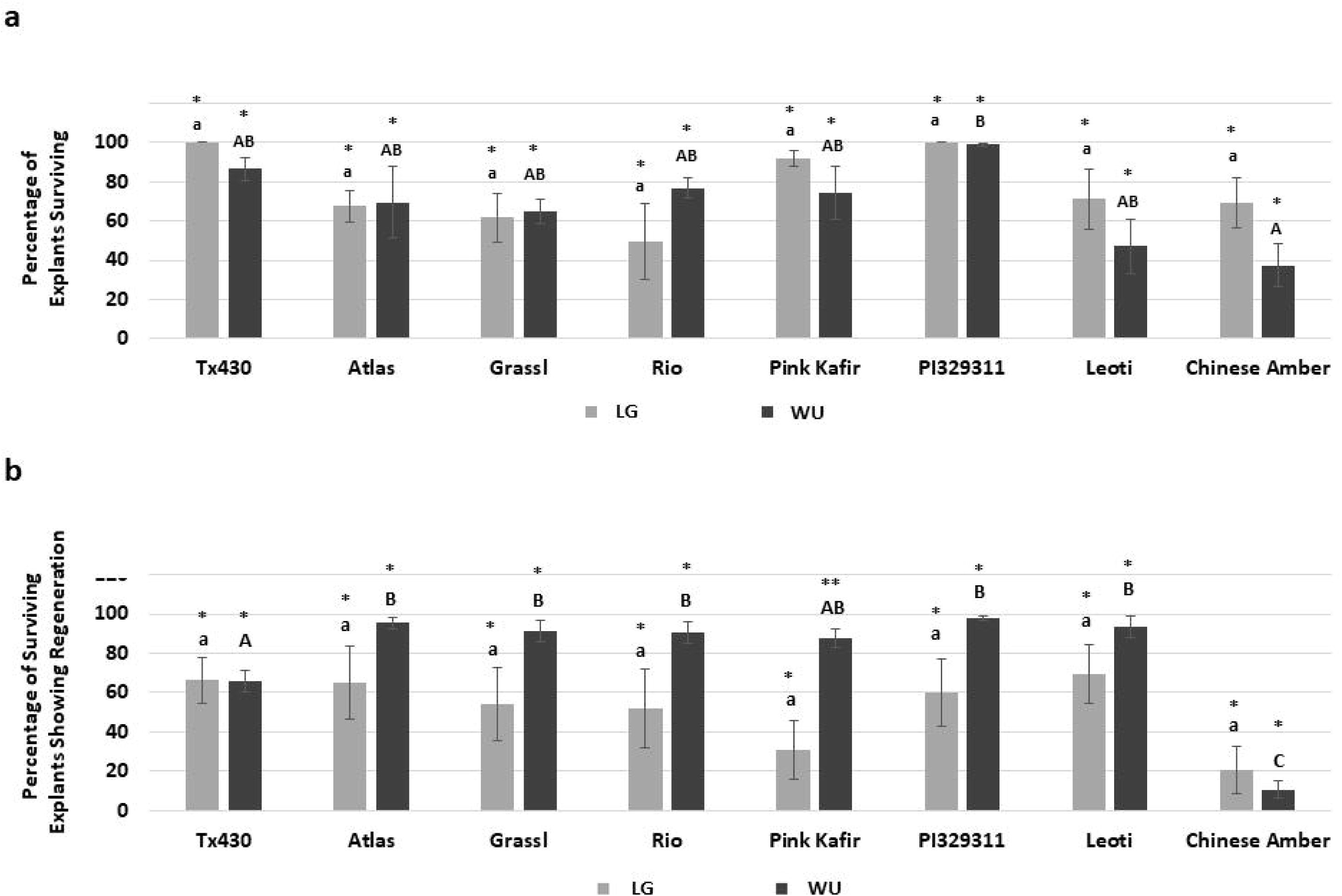
Impact of LG and WU protocols on overall explant response for the various genotypes. a) Percentage of total initial explants surviving following eight weeks in culture. Results are plotted as the mean of three independent experiments (± SE). b) Percentage of surviving explants exhibiting plantlet regeneration following eight weeks in culture. Results are plotted as the mean of three independent experiments (± SE). Data were analyzed by one-way ANOVA, and tested for significance using Tukey’s HSD Post-hoc Test. For comparisons across genotypes for the LG protocol, those with the same lowercase letter are not significantly different at P≤0.05. For comparisons across genotypes for the WU protocol, those with the same uppercase letter are not significantly different at P≤0.05. For comparisons across the two media within a genotype, those with the same asterisk pattern are not significantly different at P≤0.05.

In contrast to explant survival, some significant differences in the percentage of explants exhibiting regeneration were observed (Fig. 4b). Using the LG protocol, several genotypes (Tx430, Atlas, PI329311, Leoti) displayed an average of 60%-70% explants with regeneration. The poorest genotype response was from Chinese Amber, with an average of 21% explants displaying regeneration. Using the WU protocol, most of the genotypes (Atlas, Grassl, Rio, PI329311, Leoti) displayed an average of 91%-98% explant regeneration, and all were significantly better than Tx430 (66%). The poorest genotype response was from Chinese Amber (11%), which was significantly lower than all other genotypes using the WU protocol. We noted that all genotypes cultured using the LG protocol exhibited higher amounts of variability across experiments, while the variability was much less with the WU protocol. When comparing within genotype for both protocols, apart from genotypes Tx430 and Chinese Amber, the average percentage of explants showing regeneration was consistently greater with the WU protocol, although the only significant difference observed was with Pink Kafir.

When we determined the mean number of regenerants produced per responding explant, distinct differences were observed for genotype and media combinations (Fig. 5). All genotypes using the LG protocol were similar in regenerant production, with approximately two regenerants per responding explants, and no significant differences observed. In contrast, for the WU protocol, the mean number of regenerants was the greatest for Rio, PI329311, and Leoti, with Rio and PI329311 significantly greater than Tx430, Atlas, Grassl, Pink Kafir and Chinese Amber. In all genotypes, except for Chinese Amber, regeneration using the WU protocol was significantly better than from LG (Fig. 5). We also observed continued medium browning during this phase of plantlet development and growth. Grassl, Rio and Pink Kafir displayed more medium browning (Fig. 5b), with the degree of medium browning observed with regenerants on WU ERM either similar to, or greater than observed on LG ERM.

**Figure 5.**
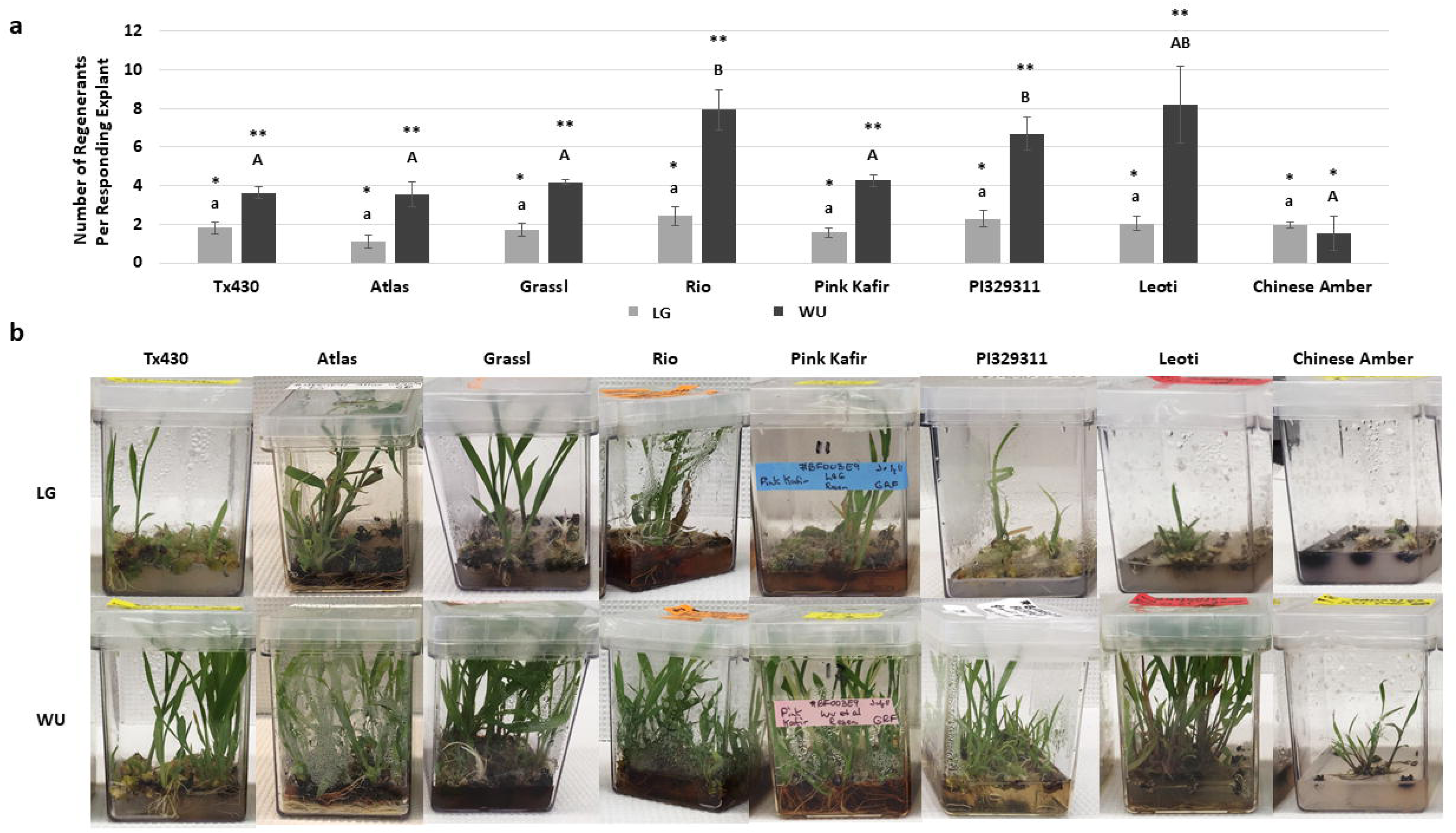
Impact of LG and WU protocols on the regeneration response for the various genotypes. a) Mean number of regenerants per responding explant following eight weeks in culture. Results are plotted as the mean of three independent experiments (± SE). b) Representative images of cultures in Magenta boxes (12 explants per box) for each genotype and protocol combination, illustrating the levels of plantlet regeneration. Data shown in (a) were analyzed by one-way ANOVA, and tested for significance using Tukey’s HSD Post-hoc Test. For comparisons across genotypes for the LG protocol, those with the same lowercase letter are not significantly different at P≤0.05. For comparisons across genotypes for the WU protocol, those with the same uppercase letter are not significantly different at P≤0.05. For comparisons across the two media within a genotype, those with the same asterisk pattern are not significantly different at P≤0.05.

During the plantlet regeneration process, we observed instances of albino plantlet development (Fig. 6), but only from three of the genotypes (Grassl, Rio, Pink Kafir). Furthermore, for these genotypes, the overwhelming majority of albino formation occurred using the WU protocol. Grassl exhibited minimal levels of albino regnerants on both LG and WU protocols, but Rio and Pink Kafir only formed albinos on the WU protocol. Pink Kafir formed albinos at a significantly higher frequency, with almost 25% of regenerating explants forming albino plantlets. Therefore, the WU protocol was much more supportive of albino formation in genotypes susceptible to this form of mutation.

**Figure 6.**
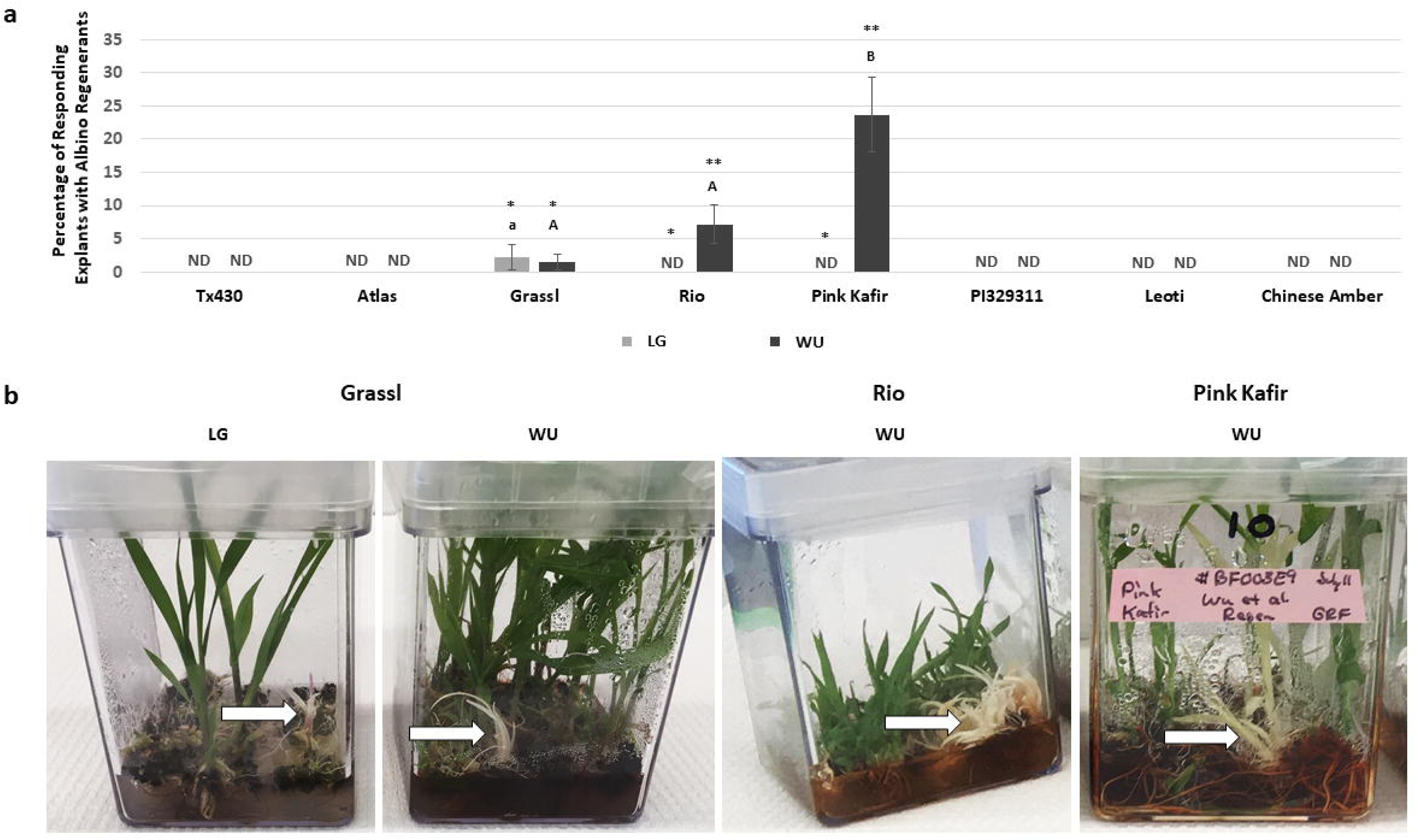
Impact of LG and WU protocols on the incidence of albino regenerants for the various genotypes. a) Percentage of regenerating explants exhibiting albino plantlet regeneration, following eight weeks in culture. Results are plotted as the mean of three independent experiments (± SE). ND, not detected. b) Representative images of cultures for genotypes exhibiting albino regeneration for specific genotype and protocol combinations. Data shown in (a) were analyzed by one-way ANOVA, and tested for significance using Tukey’s HSD Post-hoc Test. For comparisons across genotypes for the LG protocol, those with the same lowercase letter are not significantly different at P≤0.05. For comparisons across genotypes for the WU protocol, those with the same uppercase letter are not significantly different at P≤0.05. For comparisons across the two media within a genotype, those with the same asterisk pattern are not significantly different at P≤0.05.

## Discussion

Plant tissue culture responsiveness is impacted by a variety of factors, including genotype, culture medium, growth regulators and culture environment (Loyola-Vargas and Ochoa-Alejo 2018). While some plants respond easily to in vitro manipulation (e.g., carrot, tobacco), sorghum is generally considered to be a recalcitrant species in tissue culture. Hence, an understanding of the determinants of in vitro responsiveness will facilitate an expansion of this capability. In order to address this, we used a diverse subset of genotypes from the BAP (Brenton et al. 2016), which have been used as parental genotypes to create recombinant inbred line NAM populations. The ultimate goal will be to use these various populations to screen for the underlying genetic determinants based on segregation of parental traits.

Testing these NAM parental genotypes against the commonly-used grain genotype Tx430 as a reference, in combination with two different in vitro propagation protocols, allowed an assessment of genotype and protocol effects on various steps of the in vitro regeneration process. All genotypes displayed some level of response during in vitro culture with both protocols, as indicated by callus induction, callus proliferation, medium browning, explant survival and plantlet regeneration phenotypes. There were genotypic effects on the various responses, with Chinese Amber the poorest overall genotype of those tested, and most genotypes performed as well, if not better, than Tx430. Furthermore, the overall results indicated that the WU protocol was superior to LG for all genotypes tested, with the exception of Chinese Amber, indicating in vitro protocol x genotype interactions As we observed, tissue culture in sorghum, as well as other plants, is highly genotype-dependent, with varying responses observed from different genotypes (Cai et al. 1987, Elkonin et al. 1995, Kaeppler and Pedersen 1997, Liu et al. 2015, Omer et al. 2018).

The ability to predict in vitro performance of a genotype using an easily scored phenotype would be valuable. Sato et al. (2004) reported that the induction of browning during sorghum culture was a reliable indicator of poor embryogenic response. As browning is due to phenolic compound oxidation, an indicator of phenolic content might serve as a phenotype correlated with in vitro regeneration capacity. Phenolic tannins in the testa can contribute to seed coat color (Clará Valencia and Rooney 2009), suggesting that seed coat color and tannin accumulation may be potential indicators of in vitro responsiveness. Chakraborti and Ghosh (2010) reported that freshly harvested sesame seed coat color was associated with in vitro regeneration capacity, while with soybean, no relationship between embryogenic regeneration and seed coat color was noted (Ranch et al. 1985). This current work did not reveal any distinct relationship between seed coat color, tannin accumulation and regeneration response, as the three best regenerating genotypes were Rio (white seed with tannin), PI329311 (yellow seed with no tannin) and Leoti (brown seed with tannin). Sato et al (2004) also suggested that plant color (purple or tan) due to pigmentation induced under stress conditions was a marker of regenerability, with tan plants the best, and purple plants less so. In this current study, all genotypes represented purple pigmentation types, but with very distinct differences in regenerability, suggesting other significant factors are important in determining in vitro responsiveness.

The initial culture period on CIM is critical to establish the embryogenically-competent tissue needed for further differentiation and development. While all genotypes formed scutellar callus on CIM, it is possible that CIM composition differences between the two protocols had a subsequent impact on regeneration. Plant growth regulators are required for callus induction, and while both LG and WU CIM contained 1 mg/L 2,4-D as the auxin source, WU CIM also contained the cytokinin BA (0.5 mg/L). An analysis of several published sorghum studies has shown that their CIM always includes 2,4-D as the auxin source (Elkonin et al. 1995, Carvalho et al. 2004, Howe et al. 2006, Nguyen et al. 2007, Gurel et al. 2012, Chen et al. 2015, Do et al. 2016, Omer et al. 2018), sometimes combined with a cytokinin like BA (Belide et al. 2017, Espinoza-Sanchez et al. 2018) or KIN (Wernicke and Brettell 1980, Kaeppler and Pedersen 1997).

In addition, a key aspect of the WU regeneration protocol was the use of ABA. A survey of various sorghum somatic embryogenesis studies (Elkonin et al. 1995, Kaeppler and Pedersen 1997, Seetharama et al. 2000, Gurel et al. 2012, Liu and Godwin 2012, Assem et al. 2014, Wu et al. 2014, Do et al. 2016, Visarada et al. 2016, Belide et al. 2017) indicated that the inclusion of auxin(s) and cytokinin(s) are standard for regeneration, but few use ABA for embryo maturation/regeneration (Assem et al. 2014, Wu et al. 2014, Belide et al. 2017). ABA is known to promote better embryogenic responses, with higher quality embryo formation, and subsequent plantlet conversion (Merkle et al. 1995), so the superior performance of explants with the WU protocol may also reflect the inclusion of ABA during regeneration.

### Medium browning

An early response characteristic observed during the initial two-week period on CIM was the production and release of phenolic compounds, as indicated by the medium browning response. Distinct genotype and medium response profiles were noted, with some genotypes showing relatively little browning (e.g., Tx430, Atlas, Leoti), while other genotypes displayed much more browning (e.g., Grassl, Rio, Pink Kafir, PI329311, Chinese Amber). Furthermore, for some genotypes, WU CIM was much more promotive of the browning response (e.g., Tx430, Atlas, PI329311) than LG CIM. This impact of tissue culture medium on the level of browning response is attributed to the presence or absence of certain components that either serve as antioxidants to inhibit oxidative stress and phenolic oxidation or that serve as phenolic compound adsorbents to reduce their availability and toxicity (Ahmad et al. 2013, Jones and Saxena 2013). When comparing within each genotype for culture response on LG or WU CIM, the degree of browning was either similar between the two media for a genotype, or browning was greater using WU CIM; LG CIM was never more promotive of browning than WU CIM within a genotype. The higher levels of proline, as well as the incorporation of asparagine in LG CIM, may have served an antioxidant function, helping to reduce browning in those genotypes more prone to phenolic production/exudation (Szabados and Savouré 2010, Signorelli 2016). The inclusion of proline and asparagine as suggested by Elkonin et al. (1995) served to alleviate medium browning with sorghum (Carvalho et al. 2004). Studies have shown a genotype effect on browning and phenolic exudation (Cai et al. 1987, Kaeppler and Pedersen 1997), and higher browning is correlated with reduced sorghum embryogenic capacity (Kaeppler and Pedersen 1997). However, Cai et al. (1987) noted that medium modifications allowed induction of regenerable embryogenic calli from high-tannin sorghum cultivars. These results, and the results of this current study showing superior regeneration from genotypes exhibiting extensive medium browning, suggest that phenolic production and oxidation may not necessarily be prohibitive to good regeneration, and that other factors are involved. Furthermore, while the non-tannin accumulating genotype Tx430 is often used due to the lack of browning in vitro, this current study noted that both seed tannin positive and negative genotypes exhibited browning in our experiments, suggesting that browning is not specific to genotypes showing seed tannin accumulation.

### Albino regeneration

In vitro regeneration protocols have been reported as a source of somaclonal mutations during culture. We observed that three of our eight tested genotypes regenerated albino plantlets, and this was protocol dependent, with the WU protocol more prone to albino regenerant formation. The regeneration of albino plants during anther/inflorescence tissue culture has been reported for a variety of plants, including sorghum (Wen 1991; Can et al. 1998) rice (Park et al. 2013), triticale (Krzewska et al. 2015), barley (Sriskandarajah et al. 2015) and wheat (Zhao et al. 2017). In this current study, as well as that of Ma et al. (1987) and Wei and Xu (1990), albino regenerants were produced from non-reproductive tissue sorghum explants. In our study, of the three sorghum genotypes producing albino regenerants, Grassl showed the poorest response, with 2.2% of regenerating explants forming albinos when using the LG protocol, and a similar 1.5% of regenerating explants forming albinos when using the WU protocol. This was followed by Rio (7.2% of regenerating explants using the WU protocol), and Pink Kafir (23.7% of regenerating explants using the WU protocol), thus illustrating the high susceptibility of Pink Kafir to mutagenesis during in vitro development. This impact of genotype on albino regeneration has been noted during other in vitro regeneration studies (Ayed et al. 2010; Khatun et al. 2010; Park et al. 2013), as has the impact of the in vitro protocol used (Khatun et al. 2010; Park et al. 2013; Sriskandarajah et al. 2015; Dewir et al. 2018).

As explants cultured using both protocols were incubated in the same types of culture vessels and the same tissue culture chamber, our evidence of the WU protocol being more promotive of albino regeneration suggested that medium composition played a role in this response. Previous studies have shown that inclusion of casein hydrolysate reduced albinism during in vitro regeneration (Sriskandarajah et al. 2015), while the use of maltose as a carbohydrate source (Park et al. 2013), TDZ as a cytokinin source (Dewir et al. 2018) and copper supplements (Makowska et al. 2017) promoted albinism. While the WU protocol was more supportive of overall plantlet regeneration than LG, this protocol contained almost 8-fold higher levels of copper sulfate in both CIM and RM formulations, as well as the use of maltose in CIM, and TDZ in RM. Kumaravadivel and Rangasamy (1994) also noted an effect of RM growth regulator levels on albinism during plantlet regeneration. Using a sorghum anther culture system, the addition of 0.3 mg/L IAA to 2.0 mg/L BA significantly increased albino plantlet frequency. In this current study, both LG and WU RM contained IAA at 1 mg/L. However, while LG RM also contained BA, WU RM contained TDZ, zeatin and ABA. Hence, these formulation differences from LG may contribute to albinism in susceptible genotypes, and suggests that further modification to the levels of these compounds in the WU protocol could potentially reduce albinism.

Albino plantlet regeneration is associated with aberrant plastid formation and plastidic DNA deletions (Day and Ellis 1984, Dunford and Walden 1991), altered expression of photosynthesis, porphyrin and chlorophyll metabolism genes (Zhao et al. 2017), global DNA methylation changes (Duarte-Aké et al. 2016), and modified physiology (Duarte-Aké et al. 2016; Isah 2019). Furthermore, QTL studies have identified 14 chromosome regions associated with albinism in triticale (Krzewska et al. 2015), which suggested a role for oxidative stress during the proplastid to functional chloroplast transition in the promotion of albinism during in vitro regeneration. However, the underlying reasons and genetic mechanisms controlling albino plantlet formation in vitro remain to be identified. For example, how do the medium components described above interact with the genome to promote albinism? The characterization of new NAM parental lines with different susceptibility to albinism identified here under different culture conditions, as well as the availability of NAM populations derived from these various parents, provide new, additional resources to explore the genetic mechanisms underlying albino in vitro regeneration and development.

Several studies have targeted the genetic control of in vitro responsiveness in a variety of plants. Many have involved QTL identification (Bolibok and Rakoczy-Trojanowska 2006, Song et al. 2010, Tyagi et al. 2010, Trujillo-Moya et al. 2011, Yang et al. 2011, Krzewska et al. 2012, Li et al. 2013), and more recently, GWAS (Begheyn et al. 2018, Ma et al. 2018, Zhang et al. 2018). These studies have identified several genetic loci and candidate genes associated with various aspects of in vitro responsiveness. At present, the genes identified are associated with stress response regulation, cell fate change, embryogenesis and organogenesis, phytohormone metabolism and transport and chloroplast development. However, a full understanding of the suite of genes regulating in vitro responsiveness remain to be resolved, including those for sorghum.

The overall goal of this effort was to identify differential genotype-in vitro protocol responses across a variety of sorghum genotypes, in order to characterize response profiles for use in future genetic studies to identify determinants associated with in vitro regeneration. Seven NAM bioenergy sorghum genotypes and the common grain sorghum genotype Tx430 were assessed for their in vitro regeneration responses using LG and WU in vitro protocols, both previously successful with Tx430. All genotypes displayed some level of response during in vitro culture with both protocols, and distinct genotype-protocol responses were observed, with the WU protocol significantly better for plantlet regeneration. All bioenergy genotypes, with the exception of Chinese Amber, performed as well, if not better than Tx430. Genotypes displayed protocol-dependent, differential phenolic exudation responses, as indicated by medium browning. During the callus induction phase, genotypes prone to medium browning exhibited a response on WU medium which was either equal or greater than on LG medium. Genotype- and protocol-dependent albino plantlet regeneration was also noted, with three of the bioenergy genotypes showing albino plantlet regeneration, which was strongly associated with the WU protocol.

The NAM parental lines, coupled with the in vitro response characteristics described above, provide a new resource for bioenergy sorghum studies. These lines, as well as their respective recombinant inbred line populations, have been subjected to GBS, and unique molecular markers identified. The parental lines are currently in use for bioenergy (Brenton et al. 2016, Boyles et al. 2019), phytohormone (Sheflin et al. 2019), and plant-microbe interaction (Watts-Williams et al. 2019) studies. We anticipate that these parental genotypes, as well as their recombinant inbred line populations, will provide new resources and tools to identify/assess genetic loci, candidate genes, and allelic variants for their role in the regulation of in vitro responsiveness in sorghum.

## Acknowledgements

This work was supported by funds provided by the Clemson University Office of the Vice President for Research (to SK) as a component of US Department of Energy Award/Contract Number AR-0000595, as well as the Robert and Lois Coker Endowment Clemson University start-up funds (to SK).

The authors would like to thank Rick Boyles, Zach Brenton, Alex Cox, Matt Murphy and Matt Myers for their assistance with the provision of seeds, seed information, and/or the maintenance of plants in the greenhouse. The authors also thank Lindsay Shields for help with photography.

